# Illuminating the Terminal Nerve: Uncovering the Link between GnRH-1 and Olfactory Development

**DOI:** 10.1101/2023.08.31.555770

**Authors:** Enrico Amato, Ed Zandro M Taroc, Paolo E. Forni

## Abstract

During embryonic development, the olfactory placode (OP) generates migratory neurons, including olfactory pioneer neurons, cells of the terminal nerve (TN), Gonadotropin-releasing hormone-1 (GnRH-1) neurons, and other uncharacterized neurons. Pioneer neurons from the olfactory placode induce olfactory bulb morphogenesis.

In mice, GnRH-1 neurons appear in the olfactory system around mid-gestation and migrate via the terminal nerve axons to different brain regions. The GnRH-1 neurons are crucial in controlling the hypothalamic-pituitary-gonadal (HPG) axis. Kallmann syndrome is characterized by impaired olfactory system development, defective olfactory bulbs, defective secretion of GnRH-1, and infertility. The precise mechanistic link between the olfactory system and GnRH-1 development remains unclear. Studies in humans and mice highlight the importance of the Prokineticin-2/Prokineticin-Receptor-2 (Prokr2) signaling pathway in olfactory bulb morphogenesis and GnRH-1 neuronal migration. *Prokr2* loss-of-function mutations can cause Kallmann syndrome, and hence the Prokr2 signaling pathway represents a unique model to decipher the olfactory/GnRH-1 connection. We discovered that Prokr2 is expressed in the TN neurons during the critical period of GnRH-1 neuron formation, migration, and induction of olfactory bulb morphogenesis. Single-cell RNA sequencing identified that the TN is formed by neurons that are distinct from the olfactory neurons. The TN neurons express multiple genes associated with KS. Our study suggests that the aberrant development of pioneer/TN neurons might cause the KS spectrum.

**Key Points:**

1) Pioneer or terminal nerve neurons play a crucial role in initiating the development of the olfactory bulbs. We found that the Prokineticin Receptor-2 gene, associated with Kallmann syndrome, is expressed by the olfactory pioneer/terminal nerve neurons.
2) We genetically traced, isolated, and conducted Single-cell RNA sequencing on terminal nerve neurons of rodents. This analysis revealed a significant enrichment of gene expression related to Kallmann syndrome.
3) Our study indicates that the investigation of Pioneer/terminal nerve neurons should be a pivotal focal point for comprehending developmental defects affecting olfactory and GnRH-1 systems.

## Introduction

Puberty is an essential postnatal developmental process that triggers multiple hormonal changes leading to sexual and reproductive maturity (Dunkel & Quinton, 2014). Signals from the hypothalamus within the forebrain cause downstream initiation and regulation of sexual maturation and fertility (Uenoyama et al., 2019). The hypothalamic gonadotropin-releasing hormone-1 (GnRH-1) neurons (Wray, 2002) act as the main source of the GnRH-1 decapeptide that is critical in initiating the pubertal developmental cascade, maintaining sex hormone production and fertility (Schally et al., 1971).

From birds to mammals, the GnRH-1 neurons are first detectable during embryonic development in various areas of the developing olfactory pit, including respiratory, olfactory, and vomeronasal epithelium. In rodents, the GnRH-1 neurons appear to form in a region of the olfactory pit that gives rise to the vomeronasal organ (VNO) as development progresses. However, in higher species, including humans, the region of the olfactory pit where the GnRH-1 neurons form does not generate a functional VNO (Casoni et al., 2016). Nonetheless, in humans, like in mice, the GnRH-1 neurons form and migrate from the nasal area to the brain. Once in the brain, most of the GnRH-1 neurons migrate to the hypothalamus. However, several GnRH-1 cells migrate around the olfactory bulb and cortex (Casoni et al., 2016; Cho et al., 2019; Lettieri et al., 2016). Within the brain, most of the GnRH-1 neurons reside scattered in the preoptic-hypothalamic area (POA). From here, they send their axonal projections to the median eminence and into the hypophyseal-portal circulation, thus governing the hypothalamic-pituitary-gonadal axis (HPG axis) (Duittoz et al., 2021; Pitteloud et al., 2010; Tobet et al., 2001).

Disruptions in the migration of the GnRH-1 neurons, a defective release of GnRH-1, or absent/aberrant GnRH-1 signaling can lead to various forms of congenital hypogonadotropic hypogonadism (HH) (Christian et al., 1971). HH negatively affects sexual maturation, sexual function, social behavior, and fertility (Ravikumar & Crowley, 2010). HH has two main clinical categorizations, Kallmann Syndrome (KS) and normosmic idiopathic HH (niHH). In KS, there are different degrees of olfactory deficiencies spanning from a partial reduction in the sense of smell (hyposmia) to a total loss in the ability to perceive odors (anosmia) (Meschede et al., 1994). Defects in the formation of olfactory bulbs (OBs), such as OB aplasia (absence of OB) or hypoplasia (reduced OB), are common phenotypes in Kallmann patients (Martinez-Mayer & Perez-Millan, 2023). However, within families affected by KS, olfactory defects and HH do not necessarily co-segregate, suggesting that a direct link between anosmia and HH may not always exist (Balasubramanian et al., 2014). In keeping with this notion, we previously showed that in *Arx-1* null mutant mice, which cannot form olfactory bulbs (Taroc et al., 2017), the GnRH-1 neurons can still migrate into the brain, while the olfactory and vomeronasal neurons fail to connect to the brain. We revealed that the TN axons, differently from the olfactory neurons, can invade the brain with or without the olfactory bulb. This evidence suggests that the development of the OBs and the connectivity of the olfactory and vomeronasal neurons to the brain are not required for GnRH-1 neuronal migration. On this basis, we and others (Cho et al., 2019) suggested that, in mammals, the GnRH-1 neurons migrate along the axons of pioneer/ terminal nerve (TN) neurons (Demski & Schwanzel-Fukuda, 1987).

The morphogenic process leading to the formation of OBs is believed to be triggered by the interaction of early migratory olfactory pioneer neurons with the developing brain (Gong & Shipley, 1995; Wu et al., 2018). However, what the olfactory/pioneer neurons are, and their molecular characterization still needs to be addressed.

The TN is was first described in sharks (Demski, 1987), and then reported to exist in heterogeneous forms in many vertebrate species (Demski & Northcutt, 1983; Sonne et al., 2020). Interestingly, the TN has been described in cetaceans, animals that lack olfactory structures such as an olfactory bulb and olfactory epithelium (OE) (Ridgway et al., 1987), and in humans, which lack a functional VNO (Stoyanov et al., 2018). Recent data in chick suggest that the terminal nerve is a transient embryonic neuronal structure (Palaniappan et al., 2019).

The G-protein coupled receptor Prokineticin receptor 2 (Prokr2) is predominantly expressed in the mammalian central nervous system, where it has been described to regulate neurogenesis and neuronal survival (Martinez-Mayer & Perez-Millan, 2023; Ngan & Tam, 2008). Prokr2 and its ligand Prokineticin 2 (Prok2) have been implicated in neuronal migration in the developing brain and the olfactory bulb morphogenesis (Ng et al., 2005). Loss-of-function mutations in Prokr2 and its ligand, Prok2, have been linked to KS and nIHH in humans (Avbelj Stefanija et al., 2012; Balasubramanian et al., 2011; Martinez-Mayer & Perez-Millan, 2023). In line with this, mice homozygous for Prokr2 or Prok2 null mutations have severe atrophy of the reproductive system and hypoplastic olfactory bulbs. These phenotypes recapitulate the human phenotype of Kallmann syndrome (Matsumoto et al., 2006). Thus, the Prokineticin system represents a compelling candidate that may help bridge the knowledge gaps pertaining to the relevance of the terminal nerve, OB formation, and GnRH migration.

The effects of defective ProK2-Prokr2 signaling on GnRH-1 neuronal migration are likely non-cell-autonomous, as most of the GnRH-1 neurons do not express Prokr2 (Mohsen et al., 2017). Following Prokr2 expression and lineage tracing in developing rodent embryos, we identified Prokr2 expression in the early olfactory pioneer neurons, of which some appear to form the TN of mice. Through genetic lineage tracing, cell sorting, and single-cell transcriptomics, we provide evidence pointing to the fact that the terminal nerve/olfactory-pioneer neurons are distinct neuronal populations from olfactory/vomeronasal sensory neurons and GnRH-1 neurons. The critical role of pioneer neurons in triggering OB formation and the enriched expression of KS-associated genes in these cells suggests that the pathological pioneer/TN neurons’ development might be a crucial mechanistic link between anosmia and defective GnRH migration in Kallmann syndrome.

## Results

### During embryonic development, Prokr2 is expressed by populations of cells in and emerging from the developing olfactory area

We performed in-situ hybridization to follow Prokr2 expression during development (E10.5-E15.5) (Fig. 1) when the pioneer and GnRH-1 neurons form and migrate out of the developing olfactory pit. At all ages Prokr2 was found to be expressed by pioneer migratory neurons spanning between the OP to the developing forebrain (FB). Between E12.5 and E15.5, clusters of Prokr2+ cell bodies could be detected outside of the developing vomeronasal area with a pattern of expression similar to that we previously described for the putative terminal nerve marker Robo3 (Duittoz et al., 2022; Taroc et al., 2019). Notably, at these developmental stages, Prokr2 mRNA expression did not appear to be detectable in the forebrain.

**Figure 1.**
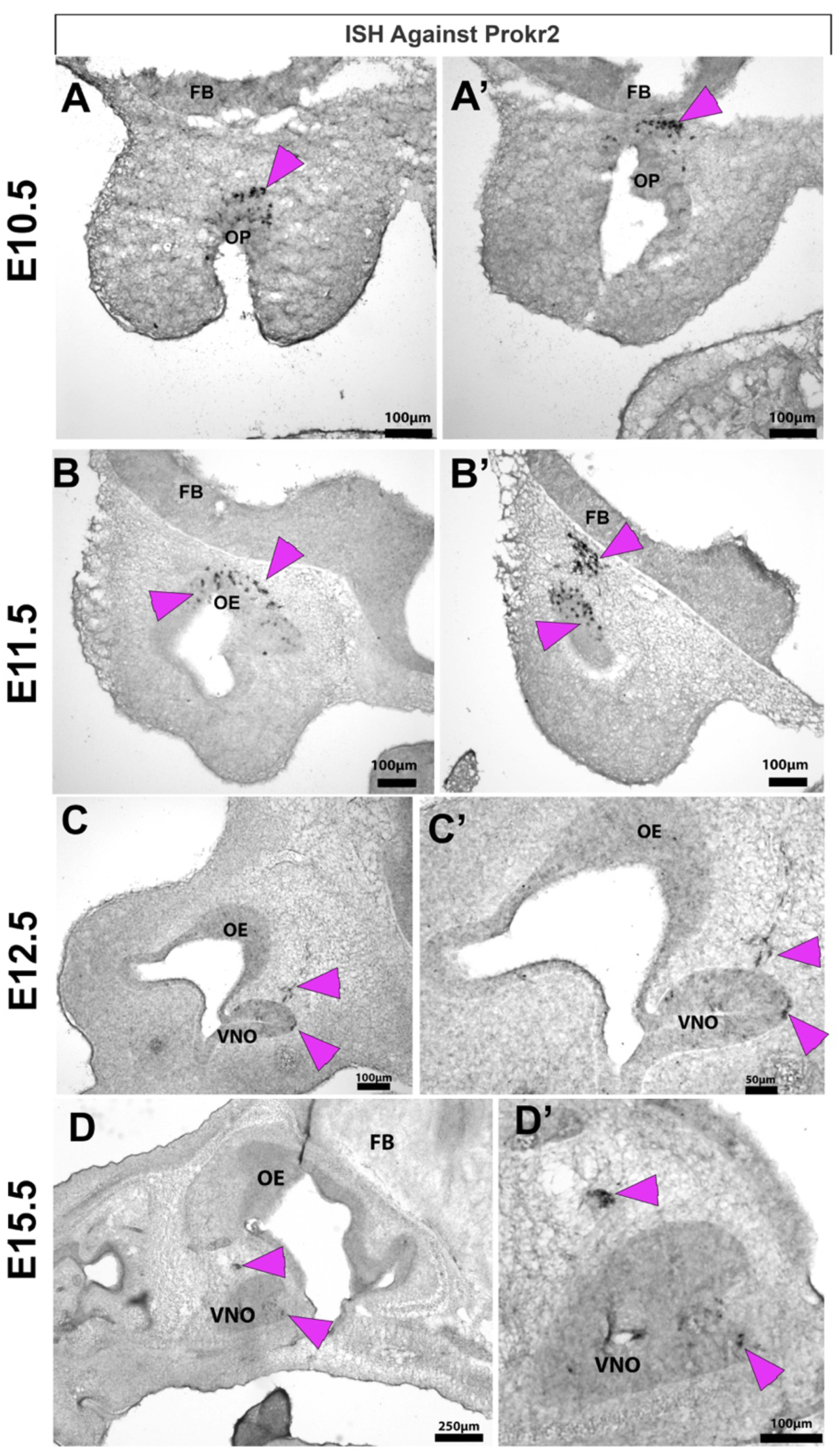
Prokr2 mRNA detected in cells of the migratory mass. **A-A ’)** In situ hybridization at E10.5, coronal section, highlights Prokr2 mRNA expression within cells (Arrowheads) near the olfactory placode (OP) and proximal to the forebrain (FB). **B-B’)** Prokr2 expressing cells at E11.5 (parasagittal) were found in the olfactory epithelium (OE) and proximal to the FB. **C-C’)** At E12.5 (parasagittal) Prokr2 is expressed in cells in and migrating out from the developing vomeronasal (VNO). **D-D’)** At E15.5, Prokr2 expression was detected in cells within the ventral portion of the developing vomeronasal organ and in clusters of cells outside of the VNO. Scale bars in A-A’, B-B’, C, D’, 100 μm; C’, 50 μm; D, 250 μm.

### Prokr2^iCre^ lineage tracing highlights the pioneer migratory neurons

Mice with null mutations for Prok2 or Prokr2 exhibit defective GnRH-1 migration; however, Prokr2 has been reported not to be expressed by the GnRH-1 neurons (Matsumoto et al., 2006). Based on this, we performed Prokr2Cre lineage tracing experiments to better understand which neuronal populations express Prokr2 between E11.5 and E14.5, stages at which the GnRH-1 neurons migrate to the brain (Gong & Shipley, 1995). In line with what we observed via in-situ hybridization, Prokr2^iCre^/R26^tdTomato^ lineage tracing at E11.5 showed recombination mostly limited to pioneer/TN neurons proximal to the developing ventral telencephalon (Fig. 2A). As suggested by the in-situ hybridization (Fig.1), at these stages no recombination was observed in the forebrain.

**Figure 2.**
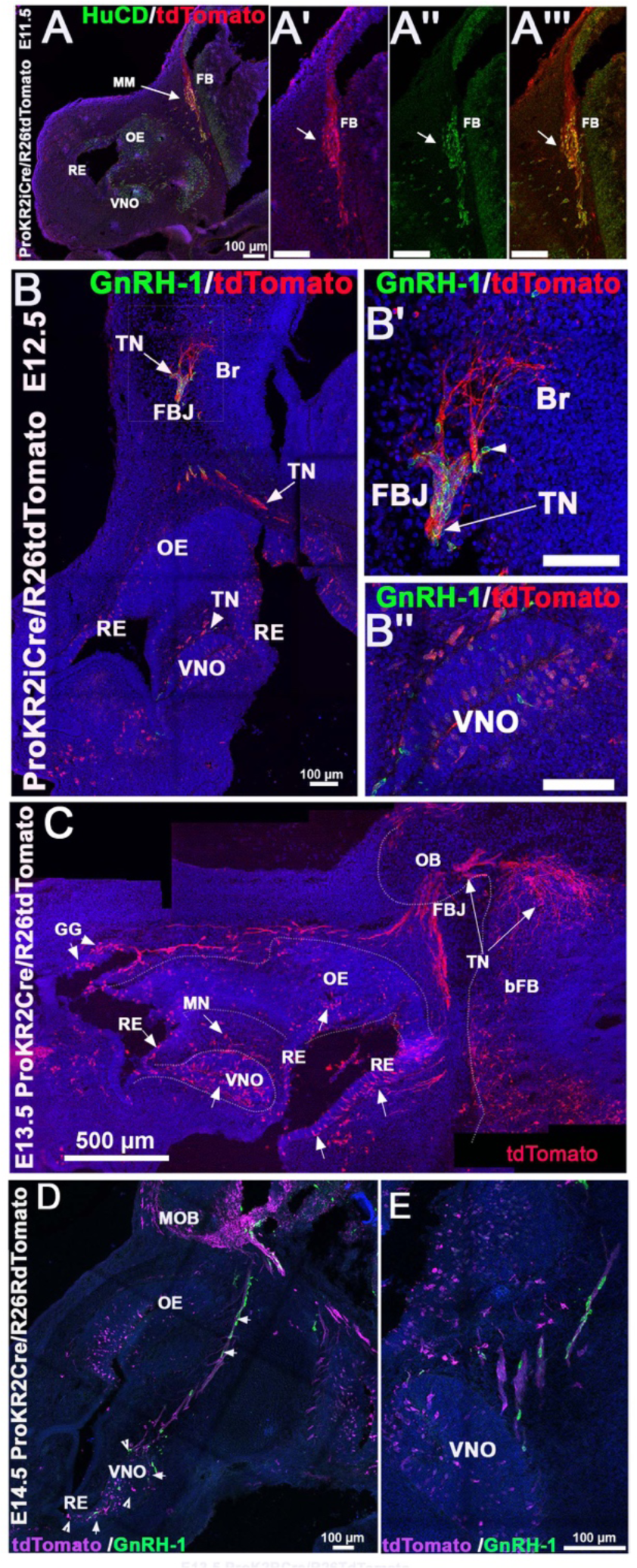
Prokr2 tracing highlights TN/pioneer neurons. **A-A ’’’)** Immunofluorescent staining of HuCD reveals a mass of Prokr2 traced neurons at E11.5. These neurons could be the putative terminal nerve or pioneer neurons of the olfactory bulb. **B-B’’)** At E12.5, GnRH-1 neurons negative for Prokr2 tracing can be seen migrating along TN axons traced for Prokr2 into the brain. Prokr2 traced neurons can be seen both within and leaving the VNO. **C)** Prokr2 traced neurons at E13.5 can be seen projecting from the nasal area toward the basal forebrain (BFB) branching in a distinct pattern. **D-E)** At E15.5 the GnRH-1 neurons (green, arrows) can be seen migrating near the traced stream of neurons (magenta, notched arrowheads) projecting from the nose to the brain. **F-F’’)** Double immunostaining of Map2 with Prokr2 tracing at E13.5 shows that both Prokr2 traced neurons within the VNO and migrating are positive for Map2 expression. **G-G’’’)** Immunostaining for Cleaved Caspase 3 on Prokr2 traced animals at E13.5 reveals that traced cells near the forebrain junction (FBJ) are positive for the cell death marker. Scale bars in A-A’’’, B-B’’, D, E 100 μm; C, 500 μm.

Between E12.5 and E13.5, the Prokr2 tracing highlighted cell bodies and axons of the putative terminal nerve/pioneer neurons (Gong & Shipley, 1995; Taroc et al., 2017) extended from the developing VNO and were found to be invading the ventral telencephalon (Fig. 2B).

Immunostaining against GnRH-1 neurons revealed that the GnRH-1 cells were associated with the ProKr2+ pioneer/terminal nerve (Fig. 2B-D). Quantifications at E13.5 revealed that the GnRH-1 neurons were, for the most part (82%) negative for the recombination but migrated along the ProKr2 positive projections. At this stage, traced axons were detected projecting to the basal forebrain (Fig. 2C). Notably, Prokr2 lineage tracing also highlighted chimeric recombination in cells within the RE (putative solitary chemosensory cells), sparse cells in the OE, and within the putative Grueneberg ganglion (GG).

Immunostaining against the microtubule-associated protein (Map2) at E13.5 revealed strong reactivity in the cells of the MM. As described for other cell types, we found expression in the cell soma and the axonal initial segments (Gumy et al., 2017; Teng et al., 2001). Notably, almost no Map2 immunoreactivity was found in the newly formed neurons of the OE and VNO.

Double immunostaining against Map2 and tdTomato on traced Prokr2cre animals highlighted that 85% of the Map2 positive cells were also positive for the tracing (Fig. 3A,B).

**Figure 3.**
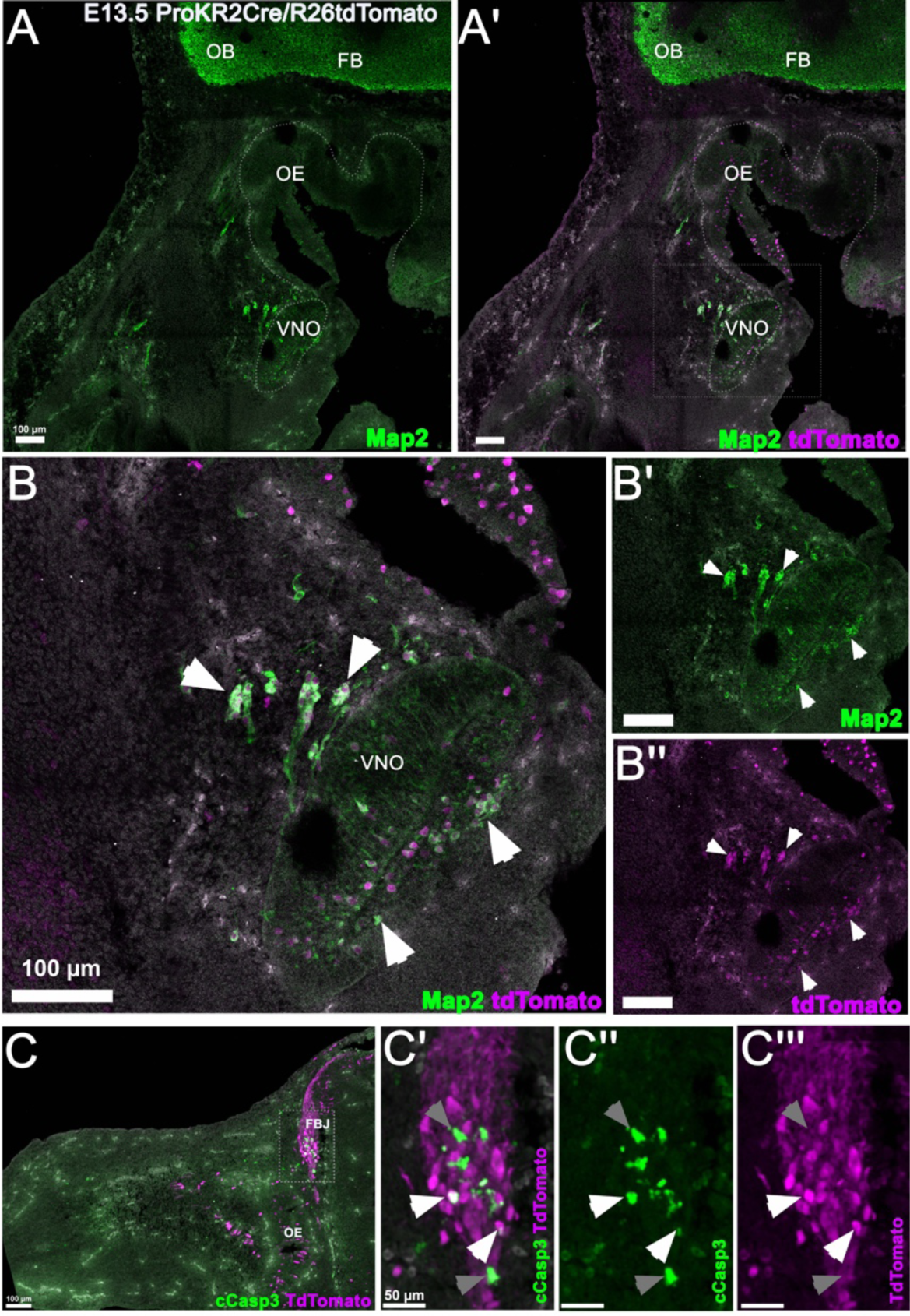
Prokr2 traced neurons express Map2 and are positive for cCasp3. **A-B ’’)** Double immunostaining of Map2 with Prokr2 tracing at E13.5 shows that both Prokr2 traced neurons within the VNO and migrating are positive for Map2 expression. Map2 immunolabelling is strong within neurons of the migratory mass and brain. Arrowheads indicate coexpression of Map2 and Prokr2 traced neurons. **C-C’’’)** Immunostaining for Cleaved Caspase 3 on Prokr2 traced animals at E13.5 reveals that many apoptotic cells are in the forebrain junction (FBJ) including Prokr2 traced-cells (grey arrows) and Prokr2 traced+ cells (white arrows). Scale bars in A-A’, B-B’’, C, 100 μm; C’-C’’’, 50 μm.

Interestingly, Map2 RNA expression and immunoreactivity of the TN/pioneer neurons appears to be more comparable to that of the forebrain neurons rather than to the one of the OSN/VSNs.

### Prokr2 Traced Cells Undergo Apoptosis Proximal to the Forebrain at E13.5

A recent report described massive apoptosis for the cells of the nasal migratory mass in chicks (Palaniappan et al., 2019). By analyzing traced animals between E13.5-E15.5, we observed detectable cleaved Caspase 3 immunoreactivity at E13.5, mostly in cells proximal to the forebrain at the putative forebrain junction (FBJ). Double immunostaining revealed that around 12% of the traced pioneer/TN cells at E13.5 were apoptotic (Fig. 3C).

These data suggest that in rodents, TN/pioneer neurons undergo cell loss proximal to the brain as described in Chick.

### Single Cell RNA Sequencing from Whole Noses

To further explore the identity of the neurons of the migratory mass/terminal nerve in the developing nasal area, we performed single-cell RNA sequencing (Sc-RNAseq) from whole embryonic mouse noses of C57 WT animals at E14.5. From these, we acquired ∼35,000 cells. Data from single cells were then grouped into clusters based on Uniform Manifold Approximation and Projections (UMAPs) (Hao et al., 2021) (Fig. 4A). Unsupervised clustering allowed us to identify nasal mesenchyme (Alx1 positive), respiratory epithelia (Krt19, Isl1 positive), and Tubb3/Ascl1/Elavl3/Foxg1/Pax6 positive clusters belonging to neurogenic regions of the olfactory pit (Fig. 4B-I). Once we identified the different cellular groups, we re-clustered the data of cells belonging to the OP (∼ 1,700 cells) without including the nasal mesenchyme (Fig. 5A). Based on gene expression enrichment, we identified respiratory epithelium (Krt14, Krt19, Tfap2a positive), stem cells (Ki-67, Hes1, Pax6, Sox2 positive), neurogenic cells (Ki-67, Ascl1 Neurog1, Neuro1), and differentiating neurons (Tubb3, Elavl3 positive) (Fig. 5C-W).

**Figure 4.**
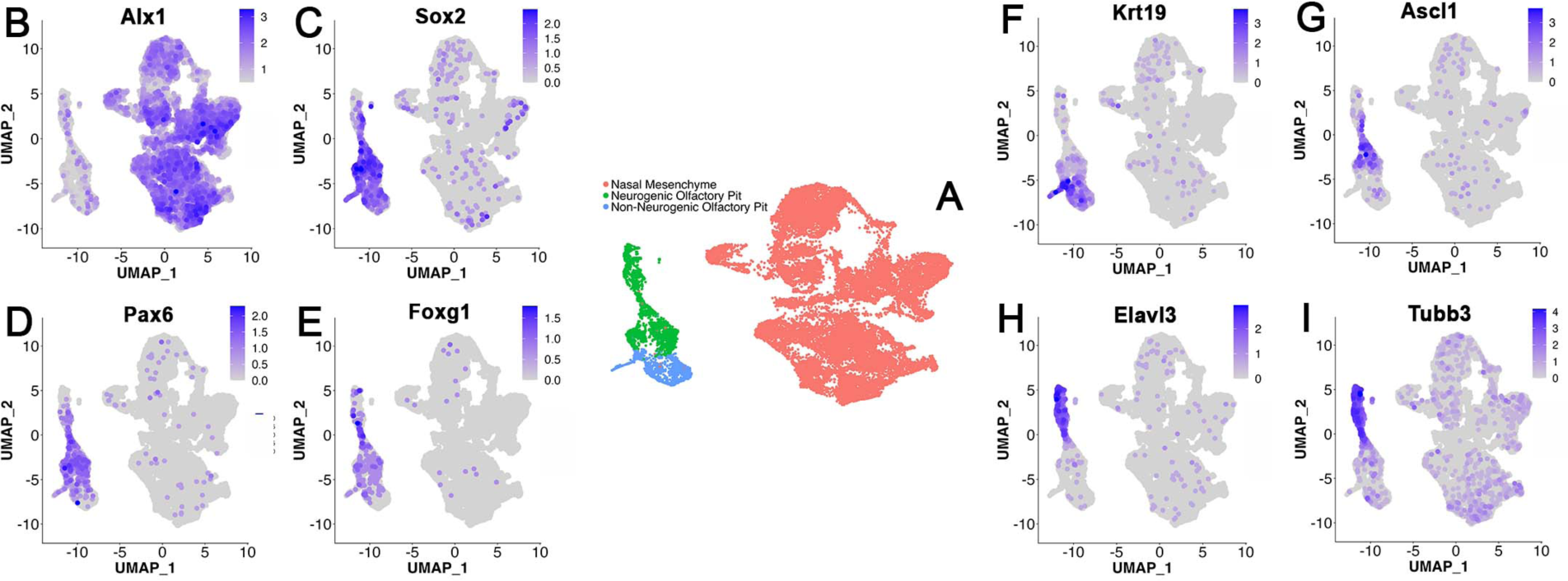
Single cell RNA-sequencing of embryonic noses. **A)** UMAP showing cell clusters isolated from E14.5 embryonic noses. A large cluster of cells were identified as mesenchyme, while a smaller, separate cluster was identified to belong to the olfactory pit, both neurogenic and non-neurogenic portions. **B-I)** Feature plots highlighting the expression of various marker genes used to group clusters into different populations of cells within the nose.

**Figure 5.**
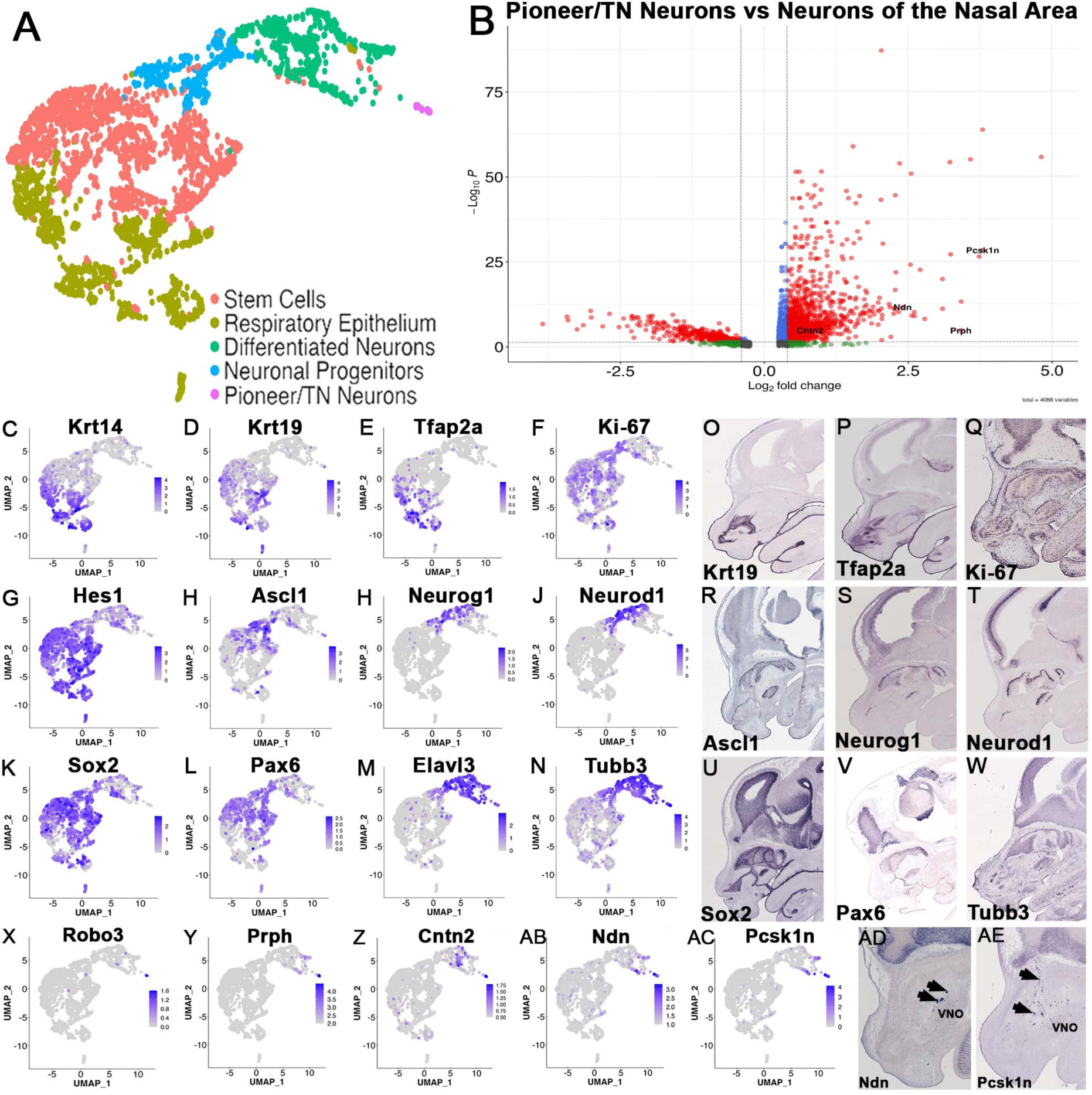
TN/pioneer neurons have a distinct transcriptome from other neurons in the nose. **A)** UMAP showing clusters present within the olfactory pit of embryonic E14.5 noses. **B)** Volcano plot showing the enriched genes of the putative migratory pioneer/TN neuron cluster (right) compared to the enriched genes of the cluster identified as differentiated neurons (left).**C-N)** Feature plots for marker genes used to identify the different regions of the olfactory area. **O-W)** In situ hybridization data at E14.5 from Gene Paint showing the expression of select genes at E14.5. **X-AC)** Feature plots showing the enriched genes within cells of the putative pioneer/TN neurons’ cluster. **AD-AE)** ISH for Ndn and Pcsk1n shows their mRNA expression in migratory cells outside the VNO.

Following the expression enrichment for previously proposed TN markers such as Robo3, Peripherin, Cntn2/Tag-1 (Fig. 5X-Z) (Casoni et al., 2012; Schwarting et al., 2004; Taroc et al., 2017; Wray et al., 1994), we also identified a small cluster of putative TN/pioneer neurons. This cluster of cells was found to be highly enriched for two genes, Proprotein Convertase Subtilisin/Kexin Type 1 Inhibitor (Pcsk1n) expression and Necdin (Ndn) (Fig. 5AB, AC). In situ hybridization showed a pattern of expression in migratory cells of the putative TN (Fig. 5AD, AE). By performing differential gene expression profiling between the putative cells of the TN and the rest of the neurons of the olfactory pit, we identified several enriched genes in our cluster of interest (Fig. 5B). This included Peripherin (Prph), Cntn2/Tag1, Pcsk1n and Ndn.

### Single Cell RNA Sequencing pioneer/TN neurons from Prokr2Cre traced embryos

To get a better understanding of the precise genetic/molecular identity of the Prokr2^iCre^/R26^tdTomato^ traced cells, we dissociated the noses of E14.5 Prokr2^iCre^/ R26^tdTomato^ embryos and performed fluorescence-activated cell sorting (FACS) to isolate the traced neurons.

We analyze 1,050 sorted cells. By integrating the Sc-RNASeq data of sorted and non-sorted nasal cells, we observed that part of the sorted cells was grouped in a cluster that formed a continuum with the previously identified unsorted putative PN/TN cluster (Fig. 6A, B).

**Figure 6.**
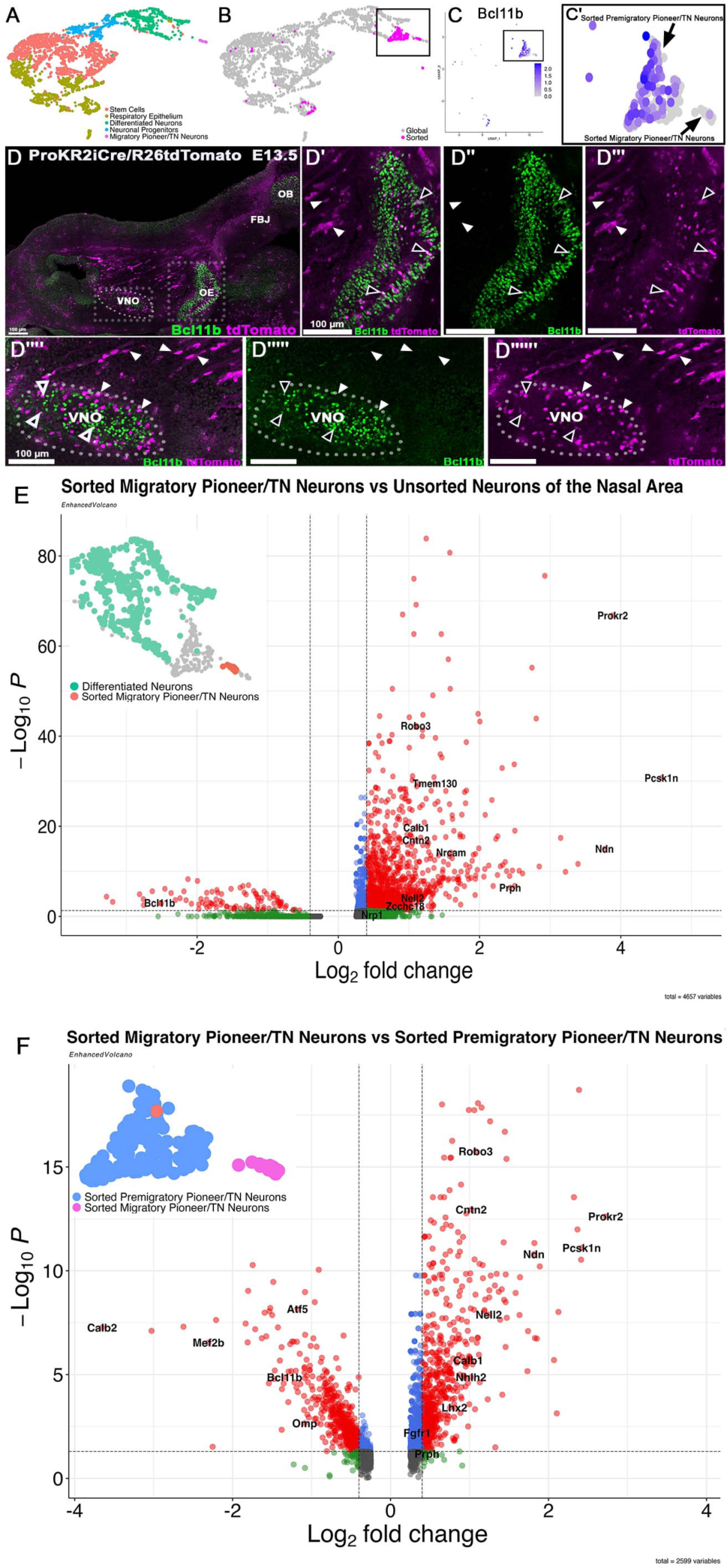
FACS sorted Prokr2^iCre^/ R26^tdTomato^ embryos have a heterogenous population of TN/pioneer neurons. **A)** UMAP of olfactory pit clusters isolated from wild-type unsorted E14.5 noses. **B)** UMAP with unsorted (grey) and sorted (magenta) cells merged. **C-D)** Feature plot of Bcl11b in the sorted cells, shown to be enriched in the sorted premigratory pioneer/TN neurons. **E-E’’’)** Bcl11b immunostaining (green) on E13.5 Prokr^iCre^ traced animals reveals that traced migratory neurons are negative for Bcl11b expression (white arrowheads) and resident pre-migratory within the VNO are positive for Bcl11b expression (black arrowheads). **F)** Volcano plot comparing enriched genes of the sorted migratory cells with all unsorted neurons. **G)** Volcano plot comparing enriched genes of the two largest sorted clusters: the putative migratory pioneer/TN neuron cluster (right) and the putative pre-migratory pioneer/TN neuron cluster (left). Scale bars in E-E’’’, 100 μm.

In addition to this, a second larger group of Prokr2^iCre^/ R26^tdTomato^ sorted cells formed a separate cluster that was found to be enriched for olfactory system markers such as OMP, Calb2 and Mef2b and Bcl11b (Jin et al., 2019; Margolis, 1972; Sammeta et al., 2007) (Fig. 6C, D, F). By performing immunolabelling against Bcl11b, we noticed that the near totality of the Prokr2^iCre^/ R26^tdTomato^ traced migratory neurons were negative for Bcl11b expression (85%), while Bcl11b expression appeared to be restricted to neurons within the developing olfactory and vomeronasal epithelia (Fig. 6D-D’’’’’’). Based on this, we classified the Prokr2^iCre^/ R26^tdTomato^ traced neurons positive for Bcl11b (Bcl11b+) as resident neurons, and the Bcl11b negative (Bcl11b-) migratory neurons.

Comparing the transcriptome of the migratory Prokr2+/TN cells to all the other neurons of the olfactory pit at E14.5, we found the enrichment of 26 genes previously linked to niHH and KS (Table 1). Notably, this list includes Sema3e and Nrp1, which are integral to the migration of the GnRH-1 neurons during embryonic development (Stamou & Georgopoulos, 2018).

**Table 1.** Genes enriched in the terminal nerve associated with Kallmann Syndrome or normosmic idiopathic hypogonadotropic hypogonadism. Known Kallmann Syndrome or normosmic idiopathic hypogonadotropic hypogonadism candidate genes enriched in the terminal nerve including their gene symbol, description, p-value, average log fold change and reference to relevant literature.

| Gene Symbol | Description | p_value | avg_log2FC | Reference |
| --- | --- | --- | --- | --- |
| Prokr2 | Prokineticin Receptor 2 | 8.05E-72 | 3.88512539 | (Matsumoto et al., 2006)<br>(Wen et al., 2019) |
| Ndn | Necdin, MAGE Family Member | 5.17E-20 | 3.76935476 | (Beneduzzi et al., 2011a) |
| Gnrh1 | Gonadotropin Releasing Hormone 1 | 5.90E-38 | 2.319258908 | (Cattanach et al., 1977)<br>(Taroc, Naik, et al., 2020) |
| Snrpn | Small Nuclear Ribonucleoprotein Polypeptide N | 2.66E-16 | 2.129445932 | (Grumbach, 2005) |
| Otx2 | Orthodenticle Homeobox 2 | 2.65E-05 | 1.471612839 | (McCabe et al., 2011) |
| Sema3e | Semaphorin 3E | 1.11E-67 | 1.458050388 | (Cariboni et al., 2015) |
| Rmst | Rhabdomyosarcoma 2 Associated Transcript | 5.76E-30 | 1.369694514 | (Stamou et al., 2020) |
| Brd2 | Bromodomain Containing 2 | 2.51E-05 | 0.627323002 | (Afonso Lopes et al., 1995)<br>(Firouzi et al., 2019) |
| Rrm2b | Ribonucleotide Reductase Regulatory TP53 Inducible Subunit M2B | 3.30E-07 | 0.576550261 | (Pitceathly et al., 2012) |
| Sos1 | SOS Ras/Rac Guanine Nucleotide Exchange Factor 1 | 2.25E-05 | 0.572023445 | (Hadziselimovic et al., 2010) |
| Lmna | Lamin A/C | 8.36E-05 | 0.539290642 | (de Andrade et al., 2020) |
| Rras | RAS Related | 2.07E-08 | 0.525486022 | (van der Kaay et al., 2016) |
| Cpe | Carboxypeptidase E | 0.00177753 | 0.517399217 | (Alsters et al., 2015) |
| Tubb3 | Tubulin Beta 3 Class III | 0.000213151 | 0.48773623 | (Chew et al., 2013)<br>(Balasubramanian et al., 2015) |
| Nrp1 | Neuropilin 1 | 2.27E-05 | 0.481371612 | (Cariboni et al., 2011) |
| Glce | Glucuronic Acid Epimerase | 3.66E-06 | 0.45641838 | (Miraoui et al., 2013) |
| Stub1 | STIP1 Homology And U-Box Containing Protein 1 | 0.006983989 | 0.426330669 | (Davis et al., 2020) |
| Rab3gap2 | RAB3 GTPase Activating Non-Catalytic Protein Subunit 2 | 0.000255373 | 0.418184055 | (Sun & Wang, 2021) |
| Mras | Muscle RAS Oncogene Homolog | 0.006374165 | 0.39834121 | (Mikec et al., 2022) |
| Aip | Aryl Hydrocarbon Receptor Interacting Protein | 0.003416918 | 0.337276059 | (Gaspar et al., 2023) |
| Las1l | LAS1 Like Ribosome Biogenesis Factor | 0.000286302 | 0.321928095 | (Gocz et al., 2022) |
| Rit1 | Ras Like Without CAAX 1 | 0.00038789 | 0.308631272 | (Deng et al., 2020) |
| Prkar2a | Protein Kinase CAMP-Dependent Type II Regulatory Subunit Alpha | 0.020431747 | 0.276596856 | (Matsumoto et al., 2006)<br>(Negri & Ferrara, 2018) |
| Ptpn11 | Protein Tyrosine Phosphatase Non-Receptor Type 11 | 0.018164779 | 0.273760812 | (Patti et al., 2023) |
| Fgf9 | Fibroblast Growth Factor 9 | 0.007879595 | 0.268214645 | (Krejci et al., 2009) |
| Magel2 | MAGE Family Member L2 | 6.18E-17 | 0.257700962 | (Mercer & Wevrick, 2009) |

### Gene expression enrichment in the Prokr2 traced pioneer/terminal nerve neurons

We and others previously described that mature and maturing GnRH-1 neurons express the transcription factor Islet1 as they migrate from the nose to the brain (Causeret et al., 2023; Palaniappan et al., 2019; Shan et al., 2020; Taroc, Katreddi, et al., 2020). By comparing the transcriptome of the sorted migratory Prokr2+ cells of the terminal nerve to the unsorted putative TN, we observed substantial differences in gene expression, suggesting the existence of two main populations of migratory neurons, the Isl1+ and Prokr2+ neurons.

The transcription factor Isl1 was significantly enriched in the Prokr2-migratory neuron cluster when compared to the migratory Prokr2+ cells. In contrast, the transcription factor Lhx2 was found to be enriched in the cells of the sorted Prokr2iCre+ migratory neurons when compared to the Isl1+ Prokr2-migratory cells (Fig. 7B).

**Figure 7.**
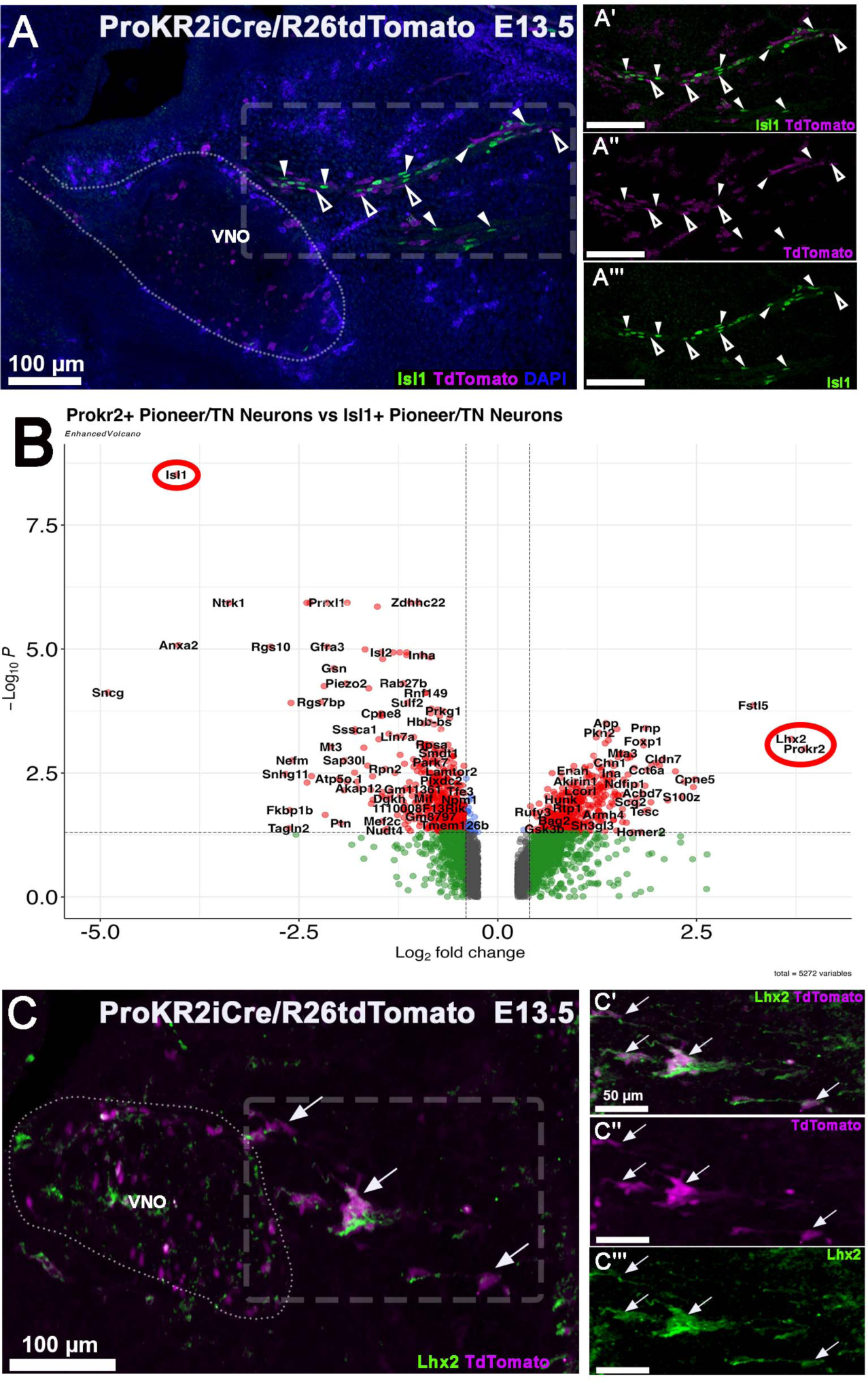
Prokr2 traced neurons express Lhx2 but not Isl1. **A-A’’’)** Prokr2cre tracing at E13.5 with Isl1 immunofluorescent staining reveals two populations of migratory cells: cells positive for Isl1 (white arrowheads) and cells negative for Isl1 that are positive for Prokr2 tracing (black arrowheads). **B)** Volcano plot comparing the enriched genes of the Prokr2+ cells (right) and the Isl1+ cells (left). **C-C’’’)** Lhx2 immunostaining on E13.5 Prokr2cre traced embryos reveals that many traced, migratory neurons express Lhx2 (white arrows). Scale bars in A-A’’’, C, 100 μm, C’-C’’’, 50 μm.

Notably, both Isl1 and Lhx2 have been recently reported to be expressed by GnRH and migratory neurons of the developing terminal nerve ganglion of chick (Palaniappan et al., 2019). However, Isl1 immunostaining on Prokr2^iCre^ traced mice visualized that the Isl1+ immunoreactivity and the Prokr2+ lineage tracing highlight two distinct migratory neuronal populations in rodents (Fig. 7A). In fact, 89% of the ProKr2 traced neurons were negative for Isl1 expression. This is in line with what we observed after GnRH and ProKR2 quantifications. Immunostaining against Lhx2 showed that many (63%) traced neurons were positive for Lhx2 expression (Fig. 7C).

### The Prokr2 cells in the vomeronasal area

In line with previous observations (Pitteloud et al., 2007) Prokr2 lineage tracing labeled neurons within the developing VNO at E13.5 and E15.5. However, at these stages, immunostaining against the VSN markers AP-2ε and Meis2 (Lin et al., 2018; Taroc, Naik, et al., 2020) showed that 90% of Prokr2 traced neurons do not express the VSNs’ markers (Fig. 8A-C). These data suggest that Prokr2 tracing highlights cell populations that are distinct from the VSNs and GnRH-1 neurons (Fig 8A,B). However, analysis at E18.5 found sparse Prokr2-traced cells within the VNO, and many of the Prokr2-traced neurons seen at previous stages were no longer present in the VNO (Fig. 8C).

**Figure 8.**
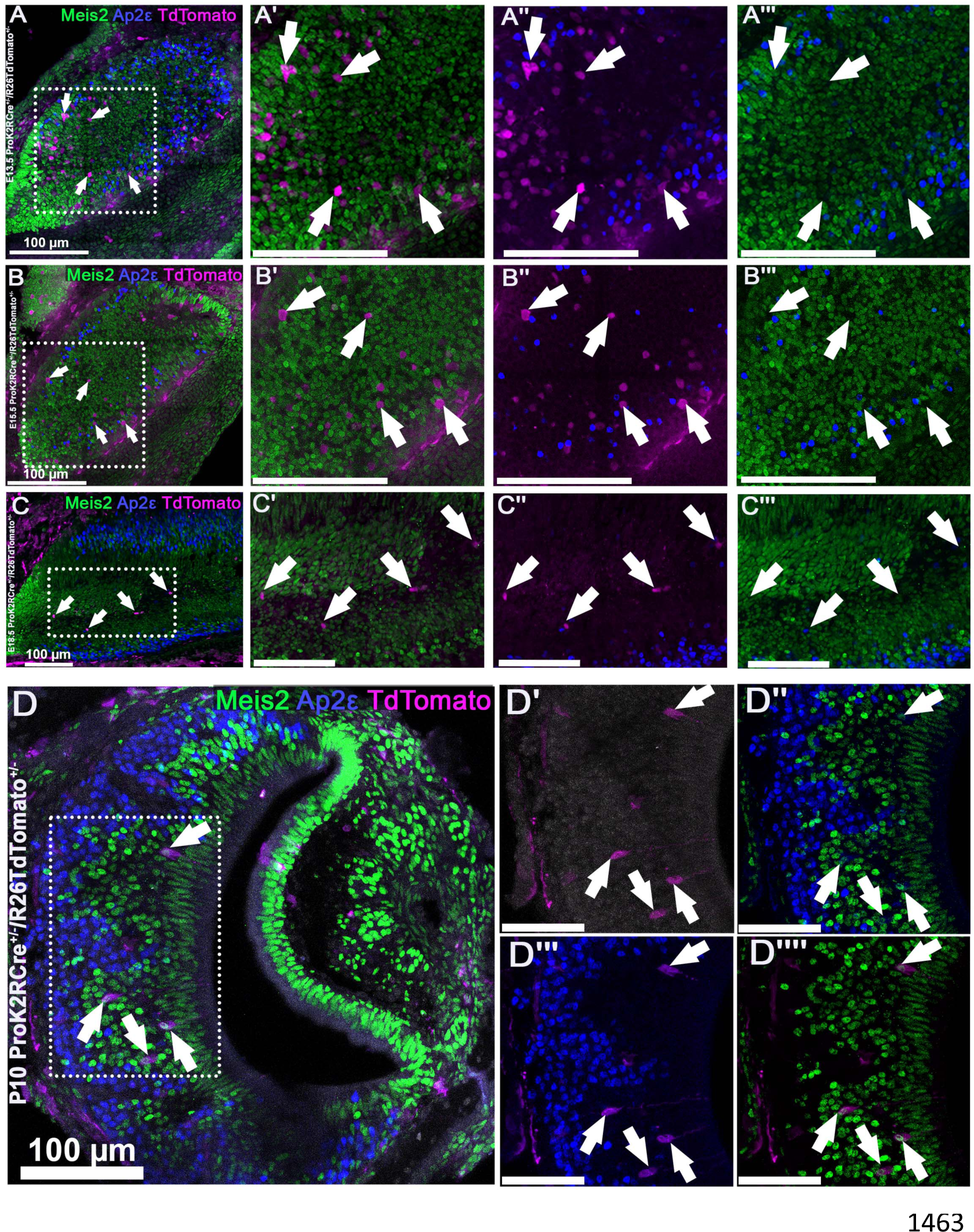
Prokr2 traced neurons are distinct from VSNs during embryonic development. **A-C)** Developmental time-course of the Prokr2 expression within the VNO from E13.5-E18.5. Immunostaining for VSN markers: AP2-ε (cyan) and Meis2 (green) against Prokr2 lineage tracing (magenta) reveals that Prokr2 traced neurons are prevalent within the VNO. Arrows indicate Prokr2 traced neurons that are negative AP2-ε and Meis2 expression. The amount of neurons that are Prokr2 traced and negative for AP2-ε and Meis2 expression within the VNO appears to decrease over time. **D-D’’’)** Prokr2iCre tracing at P10. Coronal sections of the VNO immunoassayed for AP2-ε and Meis2 reveal that the Prokr2 traced neurons present in postnatal animals are positive for the two VSN markers. Arrows indicate Prokr2 traced neurons that are positive for AP2-ε and Meis2 expression. Scale bars in A-C, 100 μm; C’-C’’’’, 50 μm.

Next, we looked at Prokr2 tracing at postnatal stage P10. Coronal sections of the VNO reveal that Prokr2 traced neurons accounted for ∼1% of the VSNs (Fig. 8D) positive for either AP-2ε or Meis2 expression. These data suggest that the majority of Prokr2 positive cells in the vomeronasal anlage are pioneer/migratory cell types that are distinct from the nonmigratory olfactory and vomeronasal sensory neurons.

Additionally, traced cells were found in the main olfactory epithelium (MOE) at P10. When paired with double immunofluorescence for odorant receptor neuron’s dorsal marker I (Nqo1) and lateral/ventral marker (Ncam2) (Bozza et al., 2009; Enomoto et al., 2019), we found that ∼95% of the traced cells were positive for either marker. These data suggest the existence of heterogeneous cell populations of Prokr2+ cells within the MOE.

### Validation of Terminal Nerve markers

To further validate the identity of the Prokr2 traced cells of the putative TN, we performed immunolabeling of Calbindin-1 (Calb1), a gene that we found to be enriched in the sorted Prokr2+ cells. Calb1 is a calcium-binding protein previously reported in migratory neurons following a similar pathway to that of the GnRH-1 neurons (Honma et al., 2004; Murakami et al., 2022). In mice, at E13.5, the GnRH-1 neurons were detected juxtaposed to Calb1 positive neurons. However, some (16%) of GnRH-1 neurons were also found to be Calb1+ (Fig. 9A-A’’).

**Figure 9.**
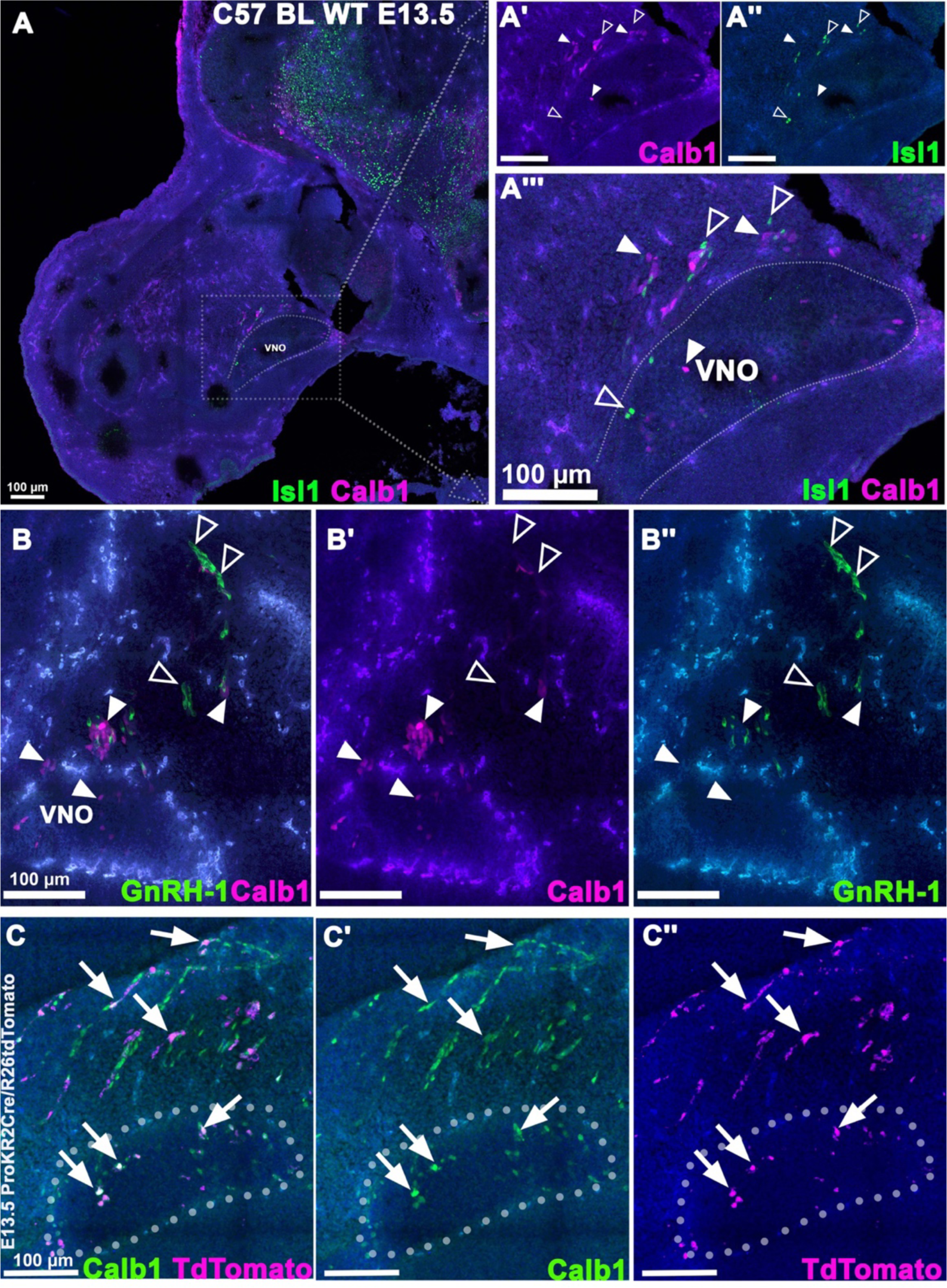
Calb1 is expressed by Prokr2 traced neurons but not Isl1 or GnRH-1 neurons. **A-A’’’)** C57 BL WT E13.5 immunofluorescent staining for Isl1 (green) and Calb1 (magenta). Both Isl1 positive (clear arrowhead) and Calb1 positive (white arrowhead) cells can be seen leaving the VNO, as two separate population of migratory cells. **B-B’’)** Immunofluorescent staining for GnRH-1 (green) shows two distinct populations of migratory neurons Isl1 positive (Including the GnRH-1 neurons) and Calb1 positive. **C-C’’’)** Prokr2iCre tracing reveals that traced neurons express Calb1 (Arrows) both in the VNO and in neurons migrating out. Scale bars in A-C, 100 μm.

In line with this, performing Calb1/Isl1 double immunostaining, we observed that 86% of the Calb1+ cells are negative for Isl1 (Fig. 9B-B’’). As suggested by Prokr2 tracing (Fig. 7) these data support various populations of migratory neurons. Calb1 immunostaining on Prokr2iCre traced embryos (Fig. 9C-C’’) found that many Calb1+ cells were traced (70%). This suggests that Calb1 can be used as a marker for Prokr2-traced neurons.

## Discussion

The olfactory placode undergoes invagination and gives rise to various populations of migratory cells (Causeret et al., 2023; Fornaro et al., 2003; Palaniappan et al., 2019; Taroc, Katreddi, et al., 2020; Valverde et al., 1993). These migratory neurons, known as pioneer neurons or neurons of the migratory mass (Fornaro et al., 2003; Maier & Gunhaga, 2009), delaminate from the placodal tissue and spread between the placode and the brain (Miller et al., 2010). In rodents, including mice, the GnRH-1 neurons, which are among the first neuronal cells to form during embryonic development, are found in the nasal migratory mass (Fornaro et al., 2003; Wray et al., 1989). The olfactory placode develops into the main olfactory system, consisting of the main olfactory epithelium and the main olfactory bulb in the brain. Some animal species also have an accessory olfactory system, comprising the vomeronasal organ and the accessory olfactory bulb, but humans lack a functional accessory olfactory system, although vestigial vomeronasal structures have been reported (Bruintjes & Bleys, 2023; Garciavelasco & Mondragon, 1991).

The prevailing model suggests that the migration of GnRH-1 neurons depends on proper olfactory development, considering the developmental defects observed in the olfactory bulbs, tracts, and sulci of Kallmann Syndrome (KS) patients (Knorr et al., 1993; Yu et al., 2022). However, evidence from mice indicates that GnRH-1 neuronal migration can occur despite a defective olfactory bulb development (Taroc et al., 2017). Interestingly, GnRH-1 neuronal migration and fertility have also been reported in human patients with arrhinia, a condition characterized by the complete absence of olfactory structures (Delaney et al., 2020).

To investigate the cellular composition of the GnRH-1 migratory track and its relation to olfactory defects in KS, researchers have explored the role of the terminal nerve or nervus terminalis, a cranial nerve that guides GnRH-1 neurons to the brain (Jennes, 1987; Jin et al., 2019; Pena-Melian et al., 2019; Schwanzel-Fukuda et al., 1989; Schwanzel-Fukuda & Pfaff, 1989; Schwanzelfukuda & Silverman, 1980; Taroc et al., 2017; Wirsigwiechmann, 1990). However, the presence of a link between the terminal nerve, GnRH-1 neuronal migration, and olfactory defects in KS still needs to be fully understood.

It is noteworthy that olfactory bulb absence or hypoplasia is common in KS patients, including those with Prokr2 and Prok2 gene mutations (Matsumoto et al., 2006; Pitteloud et al., 2007). Previous studies have shown that early pioneer neurons from the olfactory placode induce the olfactory bulb morphogenesis (Gong & Shipley, 1995; Tufo et al., 2022).

Our lineage tracing results show that Prokr2 is expressed by pioneer/terminal nerve neurons, and the migratory neurons contacting the brain before olfactory bulb formation. Moreover, in line with previous studies(Mohsen et al., 2017) most GnRH-1 neurons resulted in negative for Prokr2 but associated to axons of Prokr2-positive neurons (Fig. 2D, E).

The absence of detectable Prokr2 recombination in the GnRH-1 neurons and in the the cells forming the olfactory bulb between E11.5 and E15 (Fig. 2) strongly suggests that GnRH and olfactory bulb defects in Prokr2 null mice (Matsumoto et al., 2006; Pitteloud et al., 2007) are secondary to impaired development of pioneer/terminal nerve neurons.

Furthermore, the GnRH neurons invade the ventral forebrain before olfactory bulb formation along Prokr2-traced neuronal fibers (Fig. 2B).

Together, these data suggest that the TN migration (Fig.1,2), GnRH-1 invasion of the forebrain, and the induction of olfactory bulb (Gong & Shipley, 1995; Tufo et al., 2022) are developmentally related events. Based on this we propose that the Prokr2+ pioneer/terminal nerve neurons could have the dual function of triggering olfactory bulb formation and guiding GnRH-1 neuronal migration to the basal forebrain.

In situ hybridization experiments and lineage tracing reveled that, in the developing olfactory area, Prokr2 is expressed by migratory neurons and by limited subsets of cells in the developing vomeronasal and olfactory anlage.

Analysis of Prokr2 lineage tracing in the developing vomeronasal organ showed that most of the Prokr2-traced neurons were negative for VSN markers AP-2ε and Meis2 (Fig. 8). However, analysis at postnatal stages showed the existence of heterogeneous VSNs and OSNs positive for Prokr2 tracing.

However, our findings (Fig. 8A-D) suggest that most Prokr2+ cells originating in the olfactory placode are neurons that migrate out of the sensory epithelia as embryonic development progresses.

### Single Cell RNA Sequencing Reveals the Transcriptome of Prokr2 Cells

To better understand the cellular identity of Prokr2-expressing cells in the developing nasal area, we performed FACS sorting and single-cell sequencing of Prokr2iCre/R26tdTomato traced cells from embryonic noses. The analysis revealed that Prokr2-traced neurons were mainly distributed in two clusters (Fig. 6D), one mostly positive for Bcl11b expression and another negative for Bcl11b. Immunostaining against Bcl11b confirmed that migratory Prokr2+ pioneer/TN neurons were negative for Bcl11b, while the Prokr2-traced neurons positive for Bcl11b were within olfactory and vomeronasal epithelia. However, the postnatal analysis highlighted a minimal contribution of Prokr2+ cells to the olfactory and vomeronasal sensory lineage (Fig. 8D).

Single-cell transcriptomic analysis and immunolabelling also highlighted that the nasal migratory mass of rodents comprises two main types of migratory neurons, Prokr2+, and Isl1+ neurons.

Previous studies have shown that the Isl1 migratory neurons are, for the most part, immature or maturing GnRH-1 neurons (Shan et al., 2020; Taroc, Katreddi, et al., 2020).

The pioneer neurons, terminal nerve cells, or more vastly the so called migratory mass (MM) has been explored for many years (Demski & Schwanzel-Fukuda, 1987). Differently from what we observed in mice, previous studies in chick have indicated that the TN of chick comprises three cell types (Isl1+; GnRH1+ and Lhx2+) (Palaniappan et al., 2019). Since in mice Isl1+ cells are for the most part mature or maturing GnRH, it is possible that the discrepancies between mice and chick might reflect differences in the timing of Isl1 and GnRH expression as these migratory cells mature. In line with what described in chick we observed that the majority of the Prokr2+ cells also express Lhx2 (Fig. 7C). What the developmental and functional relationships between the Isl1 and Prokr2+ cells are needs to be further explored.

### Beyond basic developmental biology, the translational relevance

Transcriptomic analysis of the Prokr2+ Pioneer/TN cells revealed an enriched expression of the putative Grueneberg ganglia marker Pde2a (Zhi Huang, 2018), Pcsk1n, Tmem130, and Resp18. In addition to these, in the Migratory Prokr2+ cells, we identified the expression of several previously reported antigens for the putative terminal nerve/pioneer neurons, such as Calbindin1 (Fujiwara et al., 1997) (Fig. 9), Lhx2 (Palaniappan et al., 2019) (Fig. 7), Peripherin (Tobet et al., 2003), Cntn2/Tag1, Robo3. Notably, in the migratory Prokr2+ Pioneer/TN neurons, we also identified the expression of many of the candidate genes associated with KS and normosmic idiopathic hypogonadotropic hypogonadism (niHH), including Sema3e (Cariboni et al., 2007; Messina & Giacobini, 2013; Oleari et al., 2019), Prokr2, Glce (Liu & Zhi, 2022), and Necdin (ndn) (Beneduzzi et al., 2011b) (Table 1). Understanding if mutations in Kallman-related genes have a cell-autonomous effect on TN development and, therefore on GnRH-1 migration must be evaluated in the future.

Studies in Chick (Palaniappan et al., 2019) have highlighted that the TN/migratory mass undergoes massive cell death as development proceeds. By following apoptosis at E13.5 we observed similar cellular behavior in rodents (Fig. 3C). These data suggest that the TN might function as a transient developmental structure.

A technical limitation of our study is that we investigated the TN lineage by using a constitutive Prokr2Cre line that, after E14.5, does not allow us to discriminate between neurons of the TN and other brain as Prokr2Cre recombination becomes widespread in various brain regions. This limitation prevented us from further analyzing the TN connectivity or cell migration into the brain. To address this, new inducible Cre mouse models are needed to restrict Prokr2 recombination to the developmental window during TN formation.

The study provided evidence for the close relationship between the terminal nerve and GnRH-1 neurons and presented the enriched transcriptome of putative TN clusters. Lineage tracing techniques allowed the visualization of the developing TN as distinct from other neuronal populations within the nose. To fully understand TN development and its relation to KS and niHH, the identified TN markers should be further utilized in future studies.

By identifying the Prokr2+ cells as a distinct cell type from the GnRH, olfactory, and vomeronasal neurons, we set the foundation for a new chapter in the investigation of the cellular and molecular etiology of pathologies affecting olfactory and GnRH development.

Future investigations utilizing the newly discovered TN markers, identification of mutations in the TN-enriched genes, and refined mouse models will allow us to better understand the role of this nerve in normal and pathologic olfactory and sexual development.

## Materials and Methods

### Animals

The Prok2r^iCre^ mice were previously described (Mohsen et al., 2017) and ordered from the same source. R26tdTomato were maintained on a C57 BL/6J background, the mouse line (stock #007914) can be found at JAX. Prok2r^iCre^ mice were genotyped using PCR analysis from the following primers: Prokr2creF: TCCCCACGGTAGTTGTGAAG; Prokr2creR: ATTGGTTGGTGTGGTTTGCAG; Prokr2cremt: CAGCAGGTTGGAGACTTTCCT. Animals were euthanized using CO_2_, followed by cervical dislocation. Experiments used mice of either sex. All animal experiments were completed in accordance with the guidelines of the Animal Care and Use Committee (IUACUC) at the University at Albany, SUNY.

### Tissue Preparation

Embryos were collected from time-mated females where the observation of the vaginal plug was determined to be E0.5. Embryos collected were fixed in 3.7% Formaldehyde/PBS at 4°C for 2 h. Samples were then submerged in 30% sucrose overnight, then embedded and frozen in O.C.T. (Tissue-TeK) and stored at -80°C. Embedded embryos were cryosectioned using CM3050S Leica cryostat and collected on Superfrost plus slides (VWR) at 14-20 μm thickness for immunofluorescent staining.

### ISH

Digoxigenin-labeled cRNA probes were prepared via in vitro transcription (DIG RNA labeling kit; Roche Diagnostics, Basel, Switzerland) from the Prokr2 template available from Genepaint (https://gp3.mpg.de/viewer/setInfo/MH1790/12). ISH was performed as described (Forni et al., 2013) and visualized with an alkaline phosphatase conjugated anti-DIG (1:1000), and NBT/BCIP developer solution (Roche Diagnostics).

### Immunohistochemistry

Primary antibodies and dilutions used in this study: mouse (biotin)-α-HuC/D (1:1000. Invitrogen), goat-α-RFP (1:1000, Rockland), SW rabbit-α-GnRH-1 (1:6000, Susan Wray, NIH), rabbit-α-Map2 (1:200, Cell Signaling Technology), rabbit-α-Cleaved Caspase 3 (1:1000, Milipore), mouse-α-RFP (1:100, Origene), rat-α-Bcl11b (1:500, Biolegend), rabbit-α-Isl1 (1:500, Abcam), rabbit-α-Lhx2 (Bioss Antibodies, 1:250), goat-α-AP-2ε (1:500, R&D Systems), rabbit-α-Meis2 (1:1000, Abcam), mouse-α-Calb1 (1:200, Santa Cruz Biotechnology), goat-α-Nqo1 (1:700, Abcam), rabbit-α-NCAM2 (1:500, Proteintech Group). Antigen retrieval was used for experiments utilizing rabbit-α-Map2, rabbit-α-Cleaved Caspase 3, rabbit-α-Isl1, mouse-α-RFP, rat-α-Bcl11b, goat-α-AP-2ε, rabbit-α-Meis2 and mouse-α-Calb1 submerging slides in a citric acid solution heated to 95°C for 15 minutes and then cooling for 1 before continuing with standard immunofluorescence procedure. For immunofluorescent staining, secondary antibodies selected for the correct species were conjugated with Alexa Fluor-488, Alexa Flour-594, or Alexa Flour-680 (Invitrogen and Jackson Laboratories). 4′,6′-diamidino-2-phenylindole (1:3000; Sigma-Aldrich) was used to counterstain sections and coverslips were mounted using Fluorogel l (Electron Microscopy Sciences, Hatfield, PA, USA). Confocal Microscopy image acquisition was taken on a LSM 710 microscope (Zeiss, Oberkochen, Germany). Epifluorescence images were acquired on a DM4000 B LED fluorescence microscope equipped with a Leica DFC310 FX camera. Image analysis and quantifications were done using FIJI/ImageJ software. Each staining presented was replicated on three different animals.

| Antibody | Source | Dilution | Reference |
| --- | --- | --- | --- |
| Mouse (biotin) anti-HuC/D | Molecular Probes a-21272 | 1:200 | (Katreddi et al., 2022) |
| Goat anti-RFP | Rockland 200-101-379 | 1:1000 | (Chen et al., 2017) |
| SW Rabbit anti-GnRH-1 | Susan WRAY, NINDS/NIH | 1:6000 | (Wray et al., 1988) |
| Rabbit anti-Map2 | Cell Signaling Technology #4542 | 1:200 | (Li et al., 2023). |
| Rabbit anti-Cleaved Caspase 3 | Millipore AB3623 | 1:1000 | (Zhang et al., 2021) |
| Mouse anti-RFP | Origene TA180084 | 1:100 | (Katreddi et al., 2022) |
| Rat anti-Bcl11b | Biolegend 650604 | 1:500 | (Visel et al., 2004) |
| Rabbit anti-Isl1 | abcam ab20670 | 1:500 | (Qin et al., 2020) |
| Rabbit anti-Lhx2 | Bioss Antibodies bs-11200R | 1:250 | (Palaniappan et al., 2019) |
| Goat anti-AP-2ε | R&D Systems AF5060 | 1:500 | (Lin et al., 2022) |
| Rabbit anti-Meis2 | abcam ab73164 | 1:1000 | (Lin et al., 2022) |
| Mouse anti-Calb1 | Santa Cruz SC365360 | 1:200 | (Chaudhari et al., 2021) |
| Goat anti-Nqo1 | Abcam ab2346 | 1:700 | (Ihara et al., 2016) |
| Rabbit anti-NCAM2 | proteinTech Group 13850-1-AP | 1:500 | (Bozza et al., 2009) |

### Antibody Characterization

#### HuC/HuD (HuC/D)

The monoclonal HuC/D antibody was isolated from a patient with paraneoplastic encephalomyelitis with mouse as the host species. A previous study in our lab utilized this antibody for IHC (Katreddi et al., 2022), and our staining is consistent, specifically labelling neurons within the olfactory area.

#### Goat anti-red fluorescent protein (RFP)

This polyclonal antibody was derived from the mushroom polyp coral Discosoma with mouse as the host species. Western blotting labels a specific band at 27 kDA (manufacturer information). Previous work with this antibody has shown specific staining of Cre drive, tdTomato positive cells in the olfactory epithelium (Chen et al., 2017). Negative control experiments using this antibody in our lab confirm no labeling present in embryonic wild-type mouse tissue.

#### SW Gonadotropin releasing hormone-1 (GnRH-1)

This antibody was created by coupling GnRH to four different carrier proteins and using an immunization schedule on the host, rabbit, which was boosted using alternating antigen-carrier complexes. Preabsorbtion of this antibody with synthetic GnRH resulted in no immunopositive cells both in vivo and in vitro (Wray et al., 1988). Studies in our lab confirm the correct staining of the GnRH-1 neurons in the developing olfactory system as detailed in previous studies (Wray, 2010).

#### Microtubule-associated protein 2 (Map2)

This polyclonal antibody was produced by immunizing the host species, rabbit, with a synthetic peptide corresponding to carboxy terminal residues of human Map2. Map2 isoforms were confirmed to have specific bands via western blot at 280,270,70 and 75 kDa (manufacturer information). A previous study used this antibody to label Map2 in neural progenitor cells (Li et al., 2023). We confirmed Map2 expression to correlate with embryonic ISH expression data of Map2 from the online database Genepaint (Visel et al., 2004).

#### Cleaved Caspase 3

This polyclonal antibody was produced by cleavage of a specific synthetic peptide sequence with rabbit as the host species. A western blot of a non-apoptotic HeLa cell extract and an apoptotic HeLa cell extract found a specific band present at 17 kDa only for the apoptotic cell extract (manufacturer information). A recent study designed a protocol for detecting apoptotic neurons using this antibody with negative control wild type mice compared to aged mice (Zhang et al., 2021).

#### Mouse anti-red fluorescent protein (RFP)

This monoclonal antibody was produced using an RFP-monomer tagged recombinant protein expressed in E. coli with mouse as the host species. Western blotting cells transfected for the RFP-monomer were found to have a specific band at 35 kDa (manufacturer information). This antibody was used in a previous study done by our (Katreddi et al., 2022) and wild-type tissue stained for this antibody was not found to have any activity.

#### Bcl11 transcription factor B (Bcl11b)

This clone of the Bcl11b antibody has since been discontinued. This monoclonal antibody was generated from human Bcl11b with rat as the host species. From ISH expression data Bcl11b is seen to be highly enriched in the olfactory epithelium, matching the expression pattern present in our data.

#### Islet-1 (Isl1)

This polyclonal antibody was produced from a synthetic peptide corresponding to the human Islet1 amino acid sequence conjugated to keyhole limpet haemocyanin with rabbit as the host species. Similar to previous work (Qin et al., 2020) and ISH expression data (Visel et al., 2004), our stainings showed the same expression pattern of Isl1 in the brain and migratory neurons.

#### LIM homeobox 2 (Lhx2)

This polyclonal antibody was generated via conjugation of keyhole limpet haemocyanin to synthetic peptide derived from human Lhx2 with rabbit as the host species. The expression seen in our labeled tissue match with ISH expression patterns (Visel et al., 2004) and previous work in in the developing chick olfactory system (Palaniappan et al., 2019), Lhx2 is found to prevalent in the migratory mass of the nasal area.

#### Transcription factor AP-2 epsilon (AP-2ε)

This polyclonal antibody was generated from recombinant human AP-2 epsilon derived from *E. coli* with goat as the host species. Western blot of a human cell line and human placenta tissue targeted for AP-2ε both saw a specific band at 38 kDa (manufacturer information). This antibody was used in a previous paper in our lab (Lin et al., 2022) with a very strong expression pattern in the vomeronasal organ that matches with our data and ISH data.

#### Meis Homeobox 2 (Meis2)

This polyclonal antibody was generated from a synthetic peptide corresponding to human Meis2 conjugated to keyhole limpet haemocyanin with rabbit as the host species. The expression of this antibody was confirmed form similar results to a previous studies done by our lab (Lin et al., 2022) (Katreddi et al., 2022).

#### Calbindin 1 (Calb1)

This monoclonal antibody was generated from the the amino acid sequence near the N-terminus of Calb1 of human origin with mouse as the host species. Western blot of mouse brain, human kidney and rat brain tissue extracts against Calb1 showed a specific band at 28 kDa (manufacturer information). This antibody was used in a previous study to label the Purkinje cells in the cerebellar coretex (Chaudhari et al., 2021). Calb1 was found to label migratory neurons within the olfactory area matching with available ISH expression data.

#### NAD(P)H quinone dehydrogenase 1 (Nqo1)

This polyclonal antibody was generated from a synthetic peptide corresponding to the C terminal amino acid sequence of human Nqo1 with goat as the host species. Western blot of cell lysate against Nqo1 showed a single band at 30 kDa (manufacturer information). This antibody was used in previous work to label olfactory sensory neurons (Ihara et al., 2016), which matches the expression pattern in our data of the main olfactory epithelium.

#### Neural cell adhesion molecule 2 (Ncam2)

This polyclonal antibody was generated from an Ncam2 fusion protein with rabbit as the host species. Western blotting human brain tissue against Ncam2 showed a single specific band at 125 kDa (manufacturer information). Previous work has detailed Ncam2 expression within the olfactory epithelium (Bozza et al., 2009) which is what is also seen in our data.

### Single-Cell RNA Sequencing

Noses were excised from 5 CDI IGS E14.5 embryos and 1 ProKR2iCre/R26tdTomato embryo under a dissecting microscope while in sterile PBS.

Dissected noses were then further dissociated into single cells using dissociation solution. Single cells were preserved by storing and freezing them at -80 °C in (10%FBS, 90%DMSO) cell freezing media. Once both the genotype and sex of the embryos were identified, cells were defrosted quickly in a 37°C water bath, spun down, washed, and resuspended using warm 10% FBS in neurobasal medium. Cells were quantified using a T20 Automated Cell Counter (BioRad) and diluted to attain a cell concentration of ∼700-1200 cells/mL. Single-cell sequencing was performed following the 10x Chromium Next GEM Single Cell 3’ Reagent Kit v3.1 protocol using the 10x Genomics Chromium Controller within the RNA institute. Libraries generated using the 10x protocol were then submitted to be sequenced at the Center of Functional Genomics (UAlbany) using the Illumina NextSeq2000 system. The single cell suspension of the ProKR2iCre/R26tdTomato embryonic nose was sent to SingulOmics to perform FACS, generating sorted and unsorted datasets and high-throughput single-cell gene expression profiling using the 10x Genomics Chromium Platform. Data analysis was done using FASTQ files first run through the 10X CELL RANGER pipeline to align each cell’s genome to a cell-specific barcode into specific output files that can be fed into the R Package ‘Seurat’ (Satija et al., 2015). The output files were filtered through CELL RANGER, which removes barcodes that are interpreted as background noise. ‘Seurat’ was then used to do QC analysis to remove cells expressing more than 9000 genes limiting the number of dying cells analyzed; this filter was set to eliminate cells with more than 5-10% mitochondrial genes. Cell cycle regression was used to limit cell clustering based on cell cycle stage. Uniform manifold approximation and projection (UMAP) plots were used to visualize clustering. Clusters were identified for known markers for the different neuronal types such as the stem cells, neuronal progenitors, differentiated neurons and TN. Each cluster of interest went through additional QC before generating lists of enriched genes.

## Acknowledgements

We thank Dr. Andrew Poulos (University at Albany), Dr. Damian Zuloaga (University at Albany) and Dr. Ravikumar Balasubramanian (MGH, Harvard University) for their invaluable insights, comments, and critiques on this manuscript.

## Data Availability

The Single-cell RNA sequencing datasets used are available through the Gene Expression Omnibus (GEO) database, under GEO accession number GSE234871 (https://www.ncbi.nlm.nih.gov/geo/query/acc.cgi?acc=GSE234871).

